# Repeated colonization of caves leads to phenotypic convergence in catfishes (Siluriformes: *Trichomycterus*) at a small geographical scale

**DOI:** 10.1101/2020.02.25.955179

**Authors:** Juan Sebastián Flórez, Carlos Daniel Cadena, Carlos DoNascimiento, Mauricio Torres

## Abstract

Across various animal groups, adaptation to the extreme conditions of cave environments has resulted in convergent evolution of morphological, physiological, and behavioral traits. We document a Neotropical cave fish system with ample potential to study questions related to convergent adaptation to cave environments at the population level. In the karstic region of the Andes of Santander, Colombia, cave-dwelling catfishes in the genus *Trichomycterus* exhibit variable levels of reduction of eyes and body pigmentation relative to surface congeners. We tested whether cave-dwelling, eye reduced, depigmented *Trichomycterus* from separate caves in Santander were the result of a single event of cave colonization and subsequent dispersal, or of multiple colonizations to caves by surface ancestors followed by phenotypic convergence. Using mitochondrial DNA sequences to reconstruct phylogenetic relationships of *Trichomycterus* from Santander, we found that caves in this region have been colonized independently by two separate clades. Additional events of cave colonization -and possibly recolonization of surface streams- may have occurred in one of the clades, where surface and cave-dwelling populations exhibit shallow mtDNA differentiation, suggesting recent divergence or divergence in the face of gene flow. We also identified various taxonomic challenges including both a considerable number of potentially undescribed species and likely problems with the circumscription of named taxa. The system appears especially promising for studies on a wide range of ecological and evolutionary questions.

## INTRODUCTION

Animals living in caves have long been of interest for the study of adaptation by natural selection and evolutionary convergence (Juan *et al*., 2010). Across various groups –including insects in different orders, arachnids, crustaceans and several vertebrates– adaptation to the extreme conditions of cave environments has resulted in multiple instances of convergent evolution involving a variety of morphological, physiological, and behavioral traits (Jeffery, 2001; Romero, 2011; Protas *et al*., 2011; Christiansen, 2012). Some of the most striking examples of convergent evolution are found in cave-dwelling fishes of various distantly related lineages, which have consistently lost eyes and body pigmentation in comparison to their surface-living relatives (Proudlove, 2010; Borowsky, 2018; Niemiller et al. 2019).

Convergent evolution resulting from adaptation to cave environments has also been observed at the population level in fishes, with a notable example being the Mexican Tetra, *Astyanax mexicanus* (De Filippi, 1853), a well-studied taxon in which polyphyletic cave populations have reduced eyes and pigmentation, as well as unique aspects of metabolism and behavior, compared to closely related surface populations of the same species (Espinasa & Borowsky, 2001; Yoshizawa *et al*., 2010; Kowalko *et al*., 2013; Moran *et al*., 2015; Riddle *et al*., 2018). Despite their striking phenotypic differences, surface and cave populations of *A. mexicanus* are connected by gene flow, suggesting that divergence in cave specialists has been driven and maintained by selection counteracting the homogenizing effects of migration (Bradic *et al*., 2012; Herman *et al*., 2018). The Mexican Tetra has emerged as a model system to study the genetic and epigenetic basis of adaptation (Jeffery, 2001, 2009; Gross, 2012; Rohner *et al*., 2013; Gross *et al*., 2015; Gore *et al*., 2018); whether insights from the Mexican Tetra system have broad applicability to other clades adapting to caves, however, remains an open question. We here document a Neotropical cave fish system with ample potential to study questions related to adaptation to cave environments at the population level, and more broadly to test for convergence and to assess the repeatability of evolution by natural selection (Blount *et al*., 2018).

The Eastern Cordillera of the Andes of Colombia harbors a vast karst (i. e. limestone) region with hundreds of caves, most of them in the Department of Santander (Muñoz-Saba *et al*., 2013). Recent biological explorations of caves in this region have documented a wide diversity of animals restricted to cave environments, several of them endemic, including crabs, arachnids, and fishes (Campos, 2017; Villareal & García, 2016; Mesa *et al*., 2018). Among the most peculiar organisms reported in these caves are seven species of catfishes in the genus *Trichomycterus* Valenciennes, 1832, which coexist regionally with 12 surface species of the same genus occurring in open streams (Ardila-Rodriguez, 2007; 2018; Castellanos-Morales, 2018; Castellanos-Morales *et al*., 2011; Mesa *et al*., 2018; Fig. 1; Table 1). Cave-dwelling species of *Trichomycterus* show a variable degree of reduction of eyes and body pigmentation relative to surface species; some subterranean species are pigmented and have small eyes, whereas other species lack eyes entirely and are totally depigmented. In addition, the degree of eye atrophy may vary within species, among different caves (Ardila-Rodríguez, 2006; Castellanos-Morales, 2007; 2008; 2010; 2018).

**Table 1.**
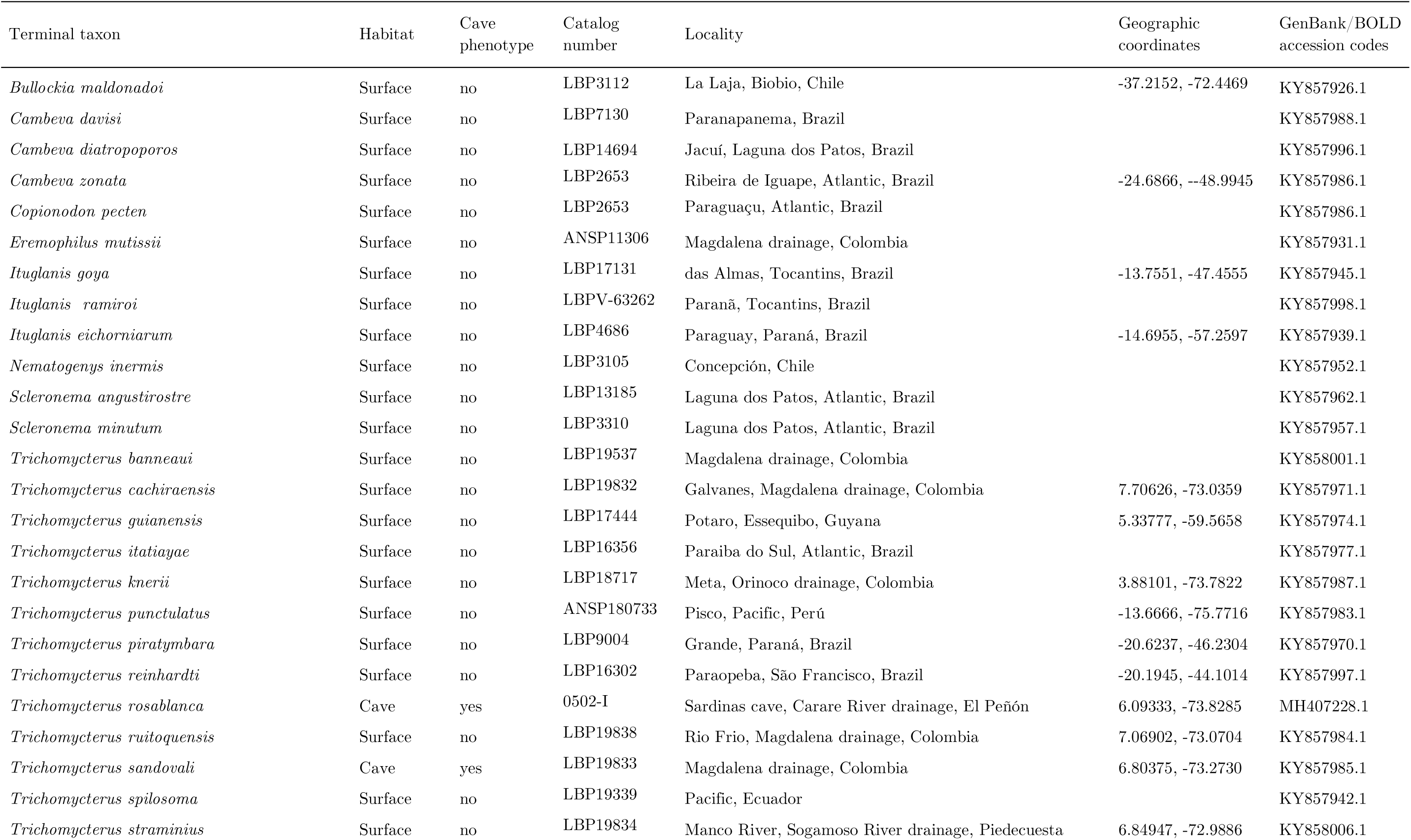
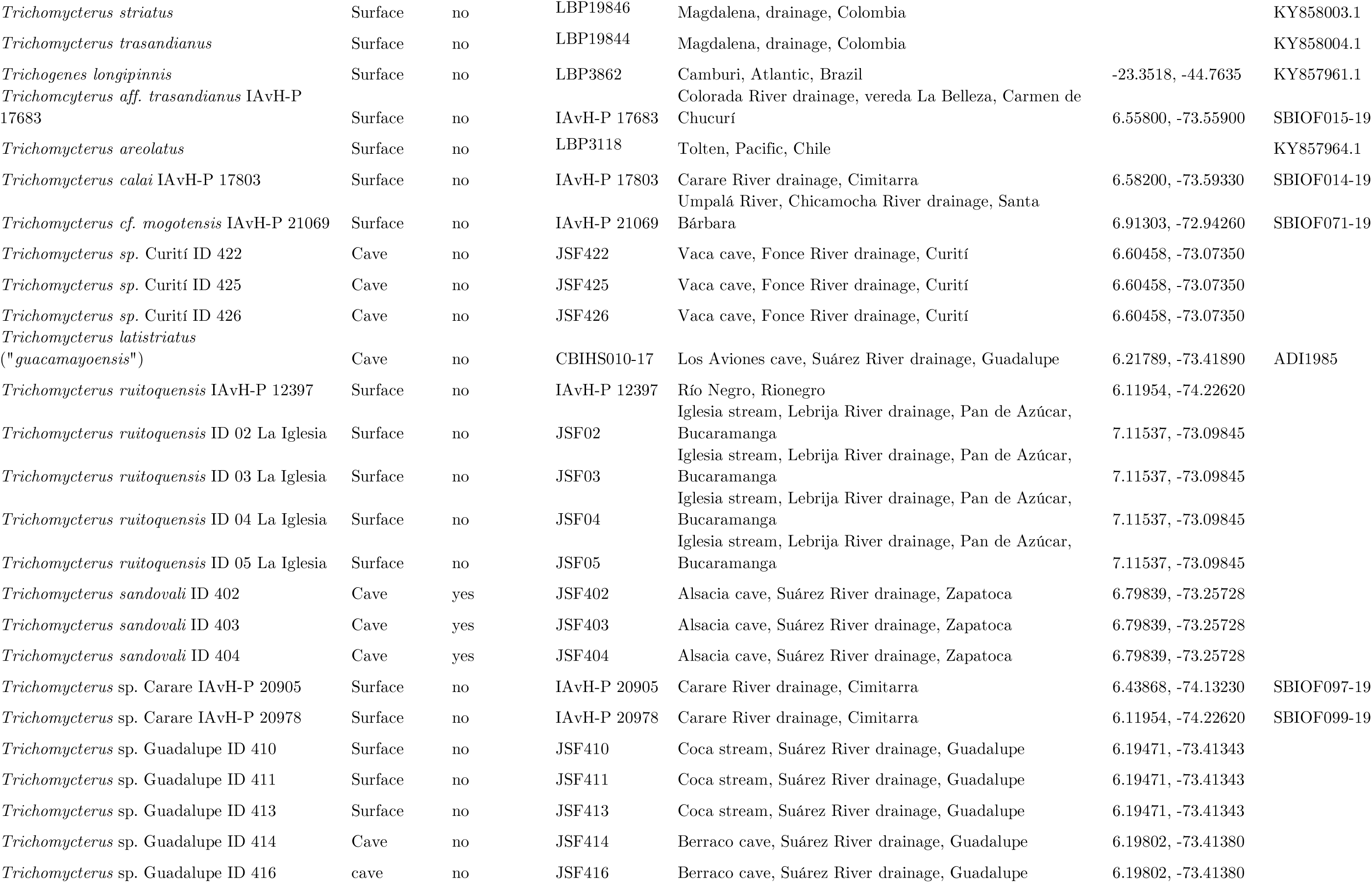
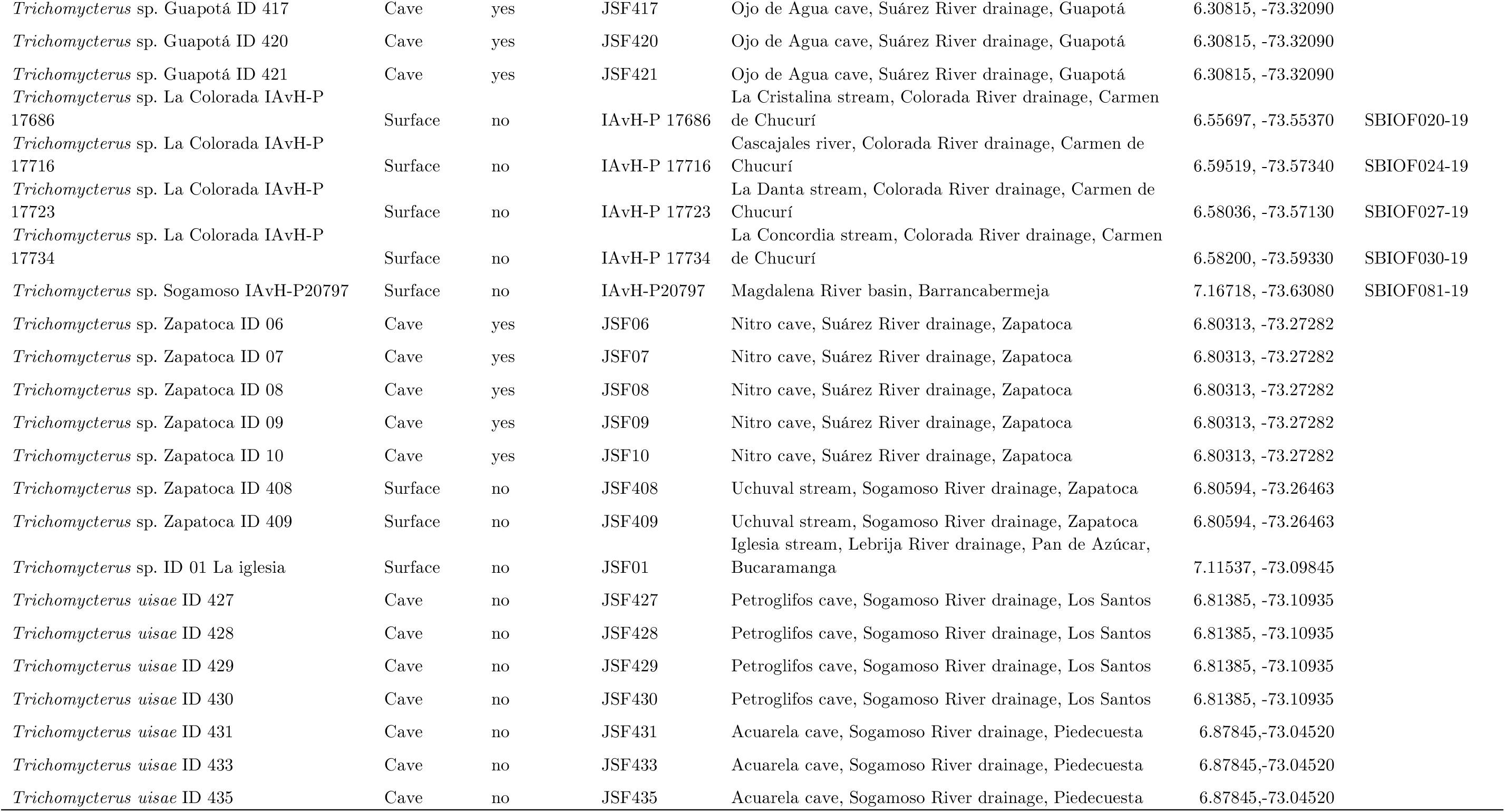
Cave and surface populations of *Trichomycterus* sampled for this study. GenBank/BOLD

| Terminal taxon | Habitat | Cave phenotype | Catalog number | Locality | Geographic coordinates | GenBank/BOLD accession codes |
| --- | --- | --- | --- | --- | --- | --- |
| *Bullockia maldonadoi* | Surface | no | LBP3112 | La Laja, Biobio, Chile | -37.2152, -72.4469 | KY857926.1 |
| *Cambeva davisi* | Surface | no | LBP7130 | Paranapanema, Brazil |  | KY857988.1 |
| *Cambeva diatropoporos* | Surface | no | LBP14694 | Jacuí, Laguna dos Patos, Brazil |  | KY857996.1 |
| *Cambeva zonata* | Surface | no | LBP2653 | Ribeira de Iguape, Atlantic, Brazil | -24.6866, -48.9945 | KY857986.1 |
| *Copionodon pecten* | Surface | no | LBP2653 | Paraguaçu, Atlantic, Brazil |  | KY857986.1 |
| *Eremophilus mutissii* | Surface | no | ANSP11306 | Magdalena drainage, Colombia |  | KY857931.1 |
| *Ituglanis goya* | Surface | no | LBP17131 | das Almas, Tocantins, Brazil | -13.7551, -47.4555 | KY857945.1 |
| *Ituglanis ramiroi* | Surface | no | LBPV-63262 | Paraná, Tocantins, Brazil |  | KY857998.1 |
| *Ituglanis eichorniarum* | Surface | no | LBP4686 | Paraguay, Paraná, Brazil | -14.6955, -57.2597 | KY857939.1 |
| *Nematogenys inermis* | Surface | no | LBP3105 | Concepción, Chile |  | KY857952.1 |
| *Scleronema angustirostre* | Surface | no | LBP13185 | Laguna dos Patos, Atlantic, Brazil |  | KY857962.1 |
| *Scleronema minutum* | Surface | no | LBP3310 | Laguna dos Patos, Atlantic, Brazil |  | KY857957.1 |
| *Trichomycterus banneaui* | Surface | no | LBP19537 | Magdalena drainage, Colombia |  | KY858001.1 |
| *Trichomycterus cachiraensis* | Surface | no | LBP19832 | Galvanes, Magdalena drainage, Colombia | 7.70626, -73.0359 | KY857971.1 |
| *Trichomycterus guianensis* | Surface | no | LBP17444 | Potaro, Essequibo, Guyana | 5.33777, -59.5658 | KY857974.1 |
| *Trichomycterus itatiyae* | Surface | no | LBP16356 | Paraíba do Sul, Atlantic, Brazil |  | KY857977.1 |
| *Trichomycterus knerii* | Surface | no | LBP18717 | Meta, Orinoco drainage, Colombia | 3.88101, -73.7822 | KY857987.1 |
| *Trichomycterus punctulatus* | Surface | no | ANSP180733 | Pisco, Pacific, Perú | -13.6666, -75.7716 | KY857983.1 |
| *Trichomycterus piratymbara* | Surface | no | LBP9004 | Grande, Paraná, Brazil | -20.6237, -46.2304 | KY857970.1 |
| *Trichomycterus reinhardti* | Surface | no | LBP16302 | Paraopeba, São Francisco, Brazil | -20.1945, -44.1014 | KY857997.1 |
| *Trichomycterus rosablanca* | Cave | yes | 0502-I | Sardinas cave, Carare River drainage, El Peñón | 6.09333, -73.8285 | MH407228.1 |
| *Trichomycterus ruitoquensis* | Surface | no | LBP19838 | Rio Frio, Magdalena drainage, Colombia | 7.06902, -73.0704 | KY857984.1 |
| *Trichomycterus sandovali* | Cave | yes | LBP19833 | Magdalena drainage, Colombia | 6.80375, -73.2730 | KY857985.1 |
| *Trichomycterus spilosoma* | Surface | no | LBP19339 | Pacific, Ecuador |  | KY857942.1 |
| *Trichomycterus stramineus* | Surface | no | LBP19834 | Manco River, Sogamoso River drainage, Piedecuesta | 6.84947, -72.9886 | KY858006.1 |
| *Trichomycterus striatus* | Surface | no | LBP19846 | Magdalena, drainage, Colombia |  | KY858003.1 |
| *Trichomycterus trasandianus* | Surface | no | LBP19844 | Magdalena, drainage, Colombia |  | KY858004.1 |
| *Trichogenes longipinnis* | Surface | no | LBP3862 | Camburi, Atlantic, Brazil | -23.3518, -44.7635 | KY857961.1 |
| *Trichomycterus aff. trasandianus* IAvH-P 17683 | Surface | no | IAvH-P 17683 | Colorada River drainage, vereda La Belleza, Carmen de Chucurí | 6.55800, -73.55900 | SBIOF015-19 |
| *Trichomycterus areolatus* | Surface | no | LBP3118 | Tolten, Pacific, Chile |  | KY857964.1 |
| *Trichomycterus calai* IAvH-P 17803 | Surface | no | IAvH-P 17803 | Carare River drainage, Cimitarra | 6.58200, -73.59330 | SBIOF014-19 |
| *Trichomycterus cf. mogotensis* IAvH-P 21069 | Surface | no | IAvH-P 21069 | Umpalá River, Chicamocha River drainage, Santa Bárbara | 6.91303, -72.94260 | SBIOF071-19 |
| *Trichomycterus sp.* Curití ID 422 | Cave | no | JSF422 | Vaca cave, Fonce River drainage, Curití | 6.60458, -73.07350 |  |
| *Trichomycterus sp.* Curití ID 425 | Cave | no | JSF425 | Vaca cave, Fonce River drainage, Curití | 6.60458, -73.07350 |  |
| *Trichomycterus sp.* Curití ID 426 | Cave | no | JSF426 | Vaca cave, Fonce River drainage, Curití | 6.60458, -73.07350 |  |
| *Trichomycterus latistriatus* ("guacamayoensis") | Cave | no | CBIHS010-17 | Los Aviones cave, Suárez River drainage, Guadalupe | 6.21789, -73.41890 | ADI1985 |
| *Trichomycterus ruitoquensis* IAvH-P 12397 | Surface | no | IAvH-P 12397 | Río Negro, Rionegro | 6.11954, -74.22620 |  |
| *Trichomycterus ruitoquensis* ID 02 La Iglesia | Surface | no | JSF02 | Iglesia stream, Lebrija River drainage, Pan de Azúcar, Bucaramanga | 7.11537, -73.09845 |  |
| *Trichomycterus ruitoquensis* ID 03 La Iglesia | Surface | no | JSF03 | Iglesia stream, Lebrija River drainage, Pan de Azúcar, Bucaramanga | 7.11537, -73.09845 |  |
| *Trichomycterus ruitoquensis* ID 04 La Iglesia | Surface | no | JSF04 | Iglesia stream, Lebrija River drainage, Pan de Azúcar, Bucaramanga | 7.11537, -73.09845 |  |
| *Trichomycterus ruitoquensis* ID 05 La Iglesia | Surface | no | JSF05 | Iglesia stream, Lebrija River drainage, Pan de Azúcar, Bucaramanga | 7.11537, -73.09845 |  |
| *Trichomycterus sandovali* ID 402 | Cave | yes | JSF402 | Alsacia cave, Suárez River drainage, Zapatoca | 6.79839, -73.25728 |  |
| *Trichomycterus sandovali* ID 403 | Cave | yes | JSF403 | Alsacia cave, Suárez River drainage, Zapatoca | 6.79839, -73.25728 |  |
| *Trichomycterus sandovali* ID 404 | Cave | yes | JSF404 | Alsacia cave, Suárez River drainage, Zapatoca | 6.79839, -73.25728 |  |
| *Trichomycterus sp.* Carare IAvH-P 20905 | Surface | no | IAvH-P 20905 | Carare River drainage, Cimitarra | 6.43868, -74.13230 | SBIOF097-19 |
| *Trichomycterus sp.* Carare IAvH-P 20978 | Surface | no | IAvH-P 20978 | Carare River drainage, Cimitarra | 6.11954, -74.22620 | SBIOF099-19 |
| *Trichomycterus sp.* Guadalupe ID 410 | Surface | no | JSF410 | Coca stream, Suárez River drainage, Guadalupe | 6.19471, -73.41343 |  |
| *Trichomycterus sp.* Guadalupe ID 411 | Surface | no | JSF411 | Coca stream, Suárez River drainage, Guadalupe | 6.19471, -73.41343 |  |
| *Trichomycterus sp.* Guadalupe ID 413 | Surface | no | JSF413 | Coca stream, Suárez River drainage, Guadalupe | 6.19471, -73.41343 |  |
| *Trichomycterus sp.* Guadalupe ID 414 | Cave | no | JSF414 | Berraco cave, Suárez River drainage, Guadalupe | 6.19802, -73.41380 |  |
| *Trichomycterus sp.* Guadalupe ID 416 | cave | no | JSF416 | Berraco cave, Suárez River drainage, Guadalupe | 6.19802, -73.41380 |  |
| *Trichomycterus* sp. Guapotá ID 417 | Cave | yes | JSF417 | Ojo de Agua cave, Suárez River drainage, Guapotá | 6.30815, -73.32090 |  |
| *Trichomycterus* sp. Guapotá ID 420 | Cave | yes | JSF420 | Ojo de Agua cave, Suárez River drainage, Guapotá | 6.30815, -73.32090 |  |
| *Trichomycterus* sp. Guapotá ID 421 | Cave | yes | JSF421 | Ojo de Agua cave, Suárez River drainage, Guapotá | 6.30815, -73.32090 |  |
| *Trichomycterus* sp. La Colorada IAvH-P 17686 | Surface | no | IAvH-P 17686 | La Cristalina stream, Colorada River drainage, Carmen de Chucurí | 6.55697, -73.55370 | SBIOF020-19 |
| *Trichomycterus* sp. La Colorada IAvH-P 17716 | Surface | no | IAvH-P 17716 | Cascajales river, Colorada River drainage, Carmen de Chucurí | 6.59519, -73.57340 | SBIOF024-19 |
| *Trichomycterus* sp. La Colorada IAvH-P 17723 | Surface | no | IAvH-P 17723 | La Danta stream, Colorada River drainage, Carmen de Chucurí | 6.58036, -73.57130 | SBIOF027-19 |
| *Trichomycterus* sp. La Colorada IAvH-P 17734 | Surface | no | IAvH-P 17734 | La Concordia stream, Colorada River drainage, Carmen de Chucurí | 6.58200, -73.59330 | SBIOF030-19 |
| *Trichomycterus* sp. Sogamoso IAvH-P20797 | Surface | no | IAvH-P20797 | Magdalena River basin, Barrancabermeja | 7.16718, -73.63080 | SBIOF081-19 |
| *Trichomycterus* sp. Zapatoca ID 06 | Cave | yes | JSF06 | Nitro cave, Suárez River drainage, Zapatoca | 6.80313, -73.27282 |  |
| *Trichomycterus* sp. Zapatoca ID 07 | Cave | yes | JSF07 | Nitro cave, Suárez River drainage, Zapatoca | 6.80313, -73.27282 |  |
| *Trichomycterus* sp. Zapatoca ID 08 | Cave | yes | JSF08 | Nitro cave, Suárez River drainage, Zapatoca | 6.80313, -73.27282 |  |
| *Trichomycterus* sp. Zapatoca ID 09 | Cave | yes | JSF09 | Nitro cave, Suárez River drainage, Zapatoca | 6.80313, -73.27282 |  |
| *Trichomycterus* sp. Zapatoca ID 10 | Cave | yes | JSF10 | Nitro cave, Suárez River drainage, Zapatoca | 6.80313, -73.27282 |  |
| *Trichomycterus* sp. Zapatoca ID 408 | Surface | no | JSF408 | Uchuval stream, Sogamoso River drainage, Zapatoca | 6.80594, -73.26463 |  |
| *Trichomycterus* sp. Zapatoca ID 409 | Surface | no | JSF409 | Uchuval stream, Sogamoso River drainage, Zapatoca | 6.80594, -73.26463 |  |
| *Trichomycterus* sp. ID 01 La iglesia | Surface | no | JSF01 | Iglesia stream, Lebrija River drainage, Pan de Azúcar, Bucaramanga | 7.11537, -73.09845 |  |
| *Trichomycterus uisae* ID 427 | Cave | no | JSF427 | Petroglifos cave, Sogamoso River drainage, Los Santos | 6.81385, -73.10935 |  |
| *Trichomycterus uisae* ID 428 | Cave | no | JSF428 | Petroglifos cave, Sogamoso River drainage, Los Santos | 6.81385, -73.10935 |  |
| *Trichomycterus uisae* ID 429 | Cave | no | JSF429 | Petroglifos cave, Sogamoso River drainage, Los Santos | 6.81385, -73.10935 |  |
| *Trichomycterus uisae* ID 430 | Cave | no | JSF430 | Petroglifos cave, Sogamoso River drainage, Los Santos | 6.81385, -73.10935 |  |
| *Trichomycterus uisae* ID 431 | Cave | no | JSF431 | Acuarela cave, Sogamoso River drainage, Piedecuesta | 6.87845, -73.04520 |  |
| *Trichomycterus uisae* ID 433 | Cave | no | JSF433 | Acuarela cave, Sogamoso River drainage, Piedecuesta | 6.87845, -73.04520 |  |
| *Trichomycterus uisae* ID 435 | Cave | no | JSF435 | Acuarela cave, Sogamoso River drainage, Piedecuesta | 6.87845, -73.04520 |  |

**Figure 1.**
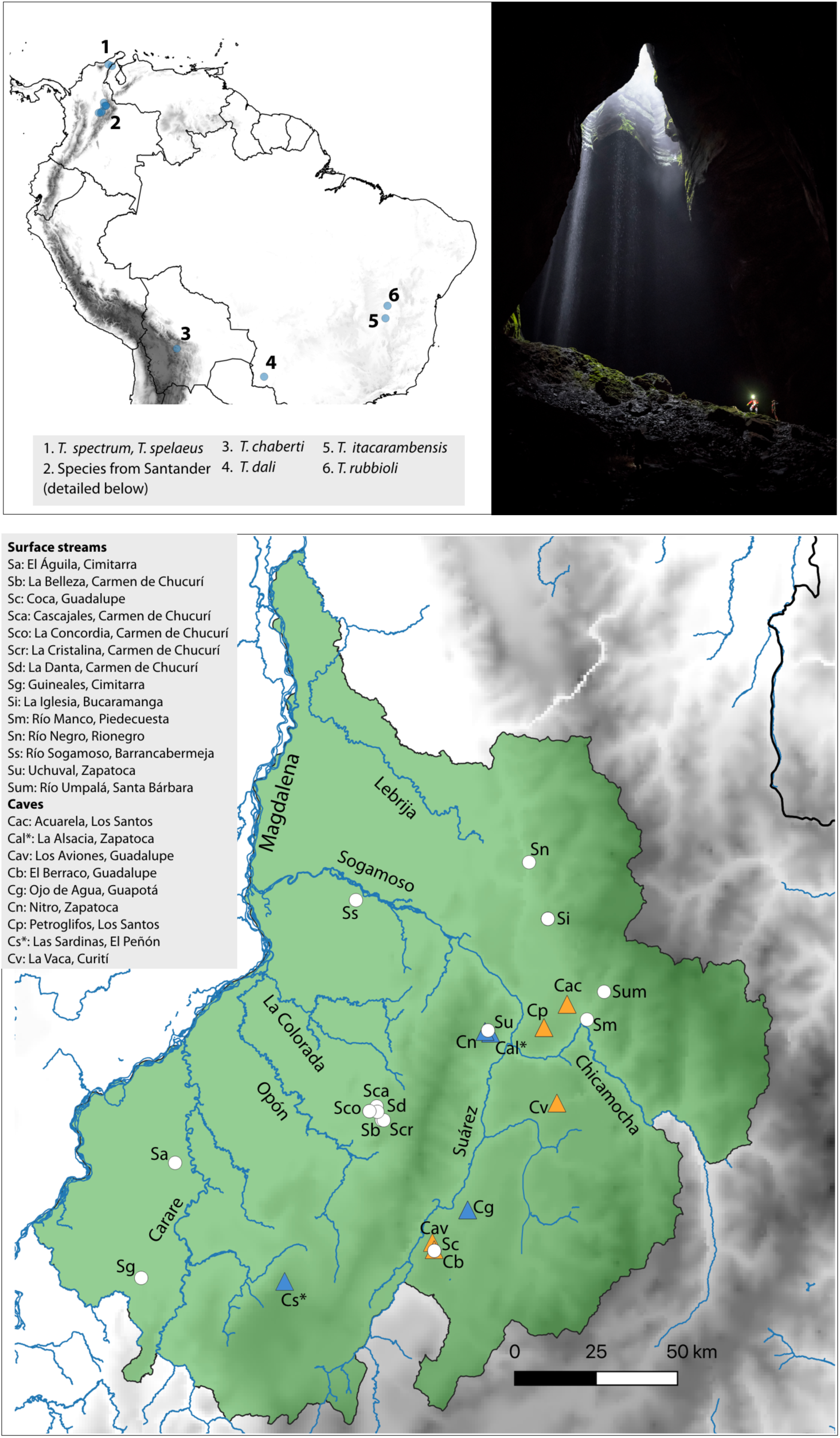
Type localities of cave species of *Trichomycterus* in the Neotropical region (top left), photograph showing an example of habitats occupied by fishes in this system (Las Sardinas Cave, the type locality of *T. rosablanca*; photo by Felipe Villegas), and map of the Department of Santander showing the collection sites of specimens of *Trichomycterus* analyzed in this study (bottom). Triangles correspond to caves, circles to surface streams. Orange triangles correspond to cave localities where fishes show reduced eyes and coloration, and blue triangles to caves where fishes have regular eyes and coloration. Cave localities are designated with letter C and surface streams with letter S. The type localities of cave species *T. rosablanca* (Cs) and *T. sandovali* (Cal) are indicated with asterisks.

Two alternative evolutionary scenarios may explain the existence of cave-dwelling *Trichomycterus* in separate caves in Santander and their phenotypic similarity involving eye atrophy and depigmentation. First, cave specialists may be the result of a single event of cave colonization, adaptation to cave environments, and subsequent subterranean vicariance or dispersal leading to their currently disjunct distributions. Alternatively, multiple colonizations to caves by surface ancestors followed by evolutionary convergence towards cave-adapted phenotypes may have occurred. We here address these alternative hypotheses using a phylogenetic approach.

Previous phylogenetic studies of the family Trichomycteridae have considered only two cave species, both from Santander: *Trichomycterus sandovali* Ardila-Rodríguez, 2006 was sampled by Ochoa *et al*. (2017, 2020) whereas *rosablanca* Mesa, Lasso, Ochoa & DoNascimiento, 2018 was included in analyses by Mesa S. *et al*. (2018). Because in those studies each cave species was independently found to be sister to the surface species *Eremophilus mutisi* Humboldt, 1805, it is possible that *T. sandovali* and *T. rosablanca* are close relatives, which would imply a single cave colonization event by *Trichomycterus* in Santander. However, available information is insufficient to reach such a conclusion owing to limited geographic and taxonomic sampling. Our field work in streams and caves of Santander and new sequence data generated from recently collected specimens from multiple localities in Colombia allows for a first robust test of hypotheses posed to explain the evolution of the group.

## MATERIAL AND METHODS

### Study System

*Trichomycterus* is one of the most taxonomically challenging groups of Neotropical fishes. The genus comprises 170 species (half the richness of its family Trichomycteridae; Fricke *et al*., 2019) ranging from Costa Rica to northern Patagonia. Species of *Trichomycterus* inhabit lowland to mountain freshwater streams, with an important fraction of its taxonomic diversity concentrated in the Colombian Andes (DoNascimiento & Prada Pedreros, 2020). To date, 12 species living in cave environments have been described from disjunct regions including Bolivia (1 species), Brazil (3), Venezuela (1), and especially the Eastern Cordillera of Colombia (Fig. 1; Bichuette & Rizzato, 2012; Ardila-Rodríguez. 2018; Castellanos-Morales, 2018; Mesa S. *et al*., 2018). *Trichomycterus* is considered a non-diagnosable, non-monophyletic taxon where species failing to meet diagnoses for other genera are often placed (Ochoa *et al*., 2017; Katz *et al*., 2018; Ochoa *et al*., 2020). One of the main issues complicating the taxonomy of *Trichomycterus* is their simplified morphology, which features relatively few characters to work with. However, a recent analysis using data from multiple genes partly clarified phylogenetic relationships in the genus, sorted species into clades largely matching large-scale geographical distributions, and made available hundreds of sequences to be employed in additional analyses (Ochoa *et al*., 2017). We here capitalize on these existing data complemented with newly generated sequences from specimens we collected to test hypotheses about evolution in cave environments in Colombia.

### Sampling

Cave and surface trichomycterids have been found along the karstic landscapes of the Eastern Cordillera of Colombia in several departments (Castellanos-Morales & Galvis, 2012; DoNascimiento *et al*., 2017). As a first approximation to this complex system, we focused our sampling efforts in the department of Santander, the region with the most extensive limestone rocks; such rocks are part of the Cumbre, Rosablanca, Paja, and Tablazo geological formations (aged from early to mid Cretaceous; Gaona-Narváez 2015; Gómez-Tapia *et al*., 2015). In Santander, meteoric and erosional processes have resulted in a karstic landscape with hundreds of caves spread in a relatively small area (∼170 km across), which remain largely unexplored (Mendoza *et al*., 2009, Muñoz-Saba *et al*., 2013).

We sampled populations of *Trichomycterus* catfishes in caves and surface streams, seeking to collect specimens from caves and adjacent streams whenever possible (Table 1). We collected 49 specimens from several populations in 2017, using hand nets or electrofishing (Samus-725 M), following methods for cave fish sampling proposed by Muñoz-Saba *et al*. (2013). Fishes were photographed, euthanized by immersion in roxicaine, and preserved in 96% ethanol. Taxonomic identification of specimens was based on comparison with original descriptions and geographic correspondence with type locality of nominal species, followed by comparative morphological examination of specimens available at the Freshwater Fish Collection of the Instituto Alexander von Humboldt (IAvH-P). For nomenclature, we followed recommendations by DoNascimiento & Prada-Pedreros (2020) regarding synonimization of some recently described taxa from Santander. All samples were deposited at IAvH-P. We also analyzed specimens collected in different drainages in Santander in 2018 during the “Santander Bio” expeditionary project (Torres & Quiñones, 2019), several of which likely correspond to undescribed species of *Trichomycterus* (Table 1). Our sampling also included *Nematogenys inermis* (Nematogenyidae), *Copionodon pecten* (Copionodontinae), and *Trichogenes longipinnis* (Trichogeninae), which we included as outgroups for phylogenetic analyses based on previous studies (Ochoa *et al*., 2017; Ochoa *et al*., 2020).

### DNA Extraction and Sequencing

We used sequences of the mitochondrial COI gene to reconstruct the phylogenetic relationships of *Trichomycterus* from Santander. This rapidly evolving marker was one of the genes used by Ochoa *et al*. (2017) to reconstruct the phylogeny of Trichomycteridae. We extracted total genomic DNA from the right pelvic fin using a phenol - chloroform - isoamyl alcohol protocol (Wasko *et al*., 2003). We amplified a 652-bp segment of the COI mitochondrial gene using primers FishF1 and FishR1 (Ward *et al*., 2005). We conducted PCRs in a volume of 25 µl, containing 2.5 µl of Buffer 10X, 1.5 µl of MgCl_2_ (50 mM), 0.5 µl of dNTPs (10 mM), 0.2 µl of Taq Polymerase (5U/µl), 1.25 µl of each primer, 1 µl of BSA (0.066 mM), 4 µl of template DNA, and 12.8 µl of ddH_2_0. The cycling included an initial denaturation at 94°C for 120 s; 34 cycles of denaturation at 94°C for 240 s, annealing at 54°C for 30 s, hybridization at 72°C for 45 s; and extension at 72°C for 600 s. PCR products were sequenced at Macrogen Inc. and the sequencing facilities of the Universidad de Los Andes (Bogotá, Colombia), resulting 41 COI gene sequences, which we visualized, edited, and aligned using the MUSCLE algorithm (Edgar, 2004) in Geneious v. 10.1.3 (Kearse *et al*., 2012). Combining our new data with existing sequences resulted in alignment comprising a total of 74 sequences (Table 1).

### Phylogenetic Analysis

We used Bayesian inference as implemented in MrBayes v. 3.2.6 (Huelsenbeck & Ronquist, 2001) and maximum likelihood in RAxML v. 8.2.10 (Stamatakis, 2014) to infer gene trees based on COI sequences. Bayesian analyses were conducted under the GTR+I+G model of nucleotide substitution, identified as the best-fit to the COI data as per the Akaike Information Criterion in PartitionFinder v. 2.1.1 (Lanfear *et al*., 2012). We ran 30 million generations, with two runs of four independent MCMC chains (three heated, one cold), sampling trees every 1000 generations, with 25% of generations discarded as burn-in. The maximum-likelihood analysis was performed under the GTR+GAMMA model and bootstrap resampling was applied to assess nodal support using the autoMRE criterion. To further examine relationships among COI haplotypes, we constructed a median-joining haplotype network using PopArt v. 1.7 (Leigh & Bryant, 2015) and redrew it manually. Finally, we asked whether a phylogeny enforcing the monophyly of cave species was less likely than unconstrained topologies using a Shimodaira–Hasegawa test (Shimodaira, 2002), implemented in the *phangorn* package (Schliep, 2011) for R (R Development Core Team, 2018).

## RESULTS

We analyzed 522 base pairs of the COI gene, of which 162 were variable (22 singletons, 140 parsimony-informative sites). Bayesian and maximum-likelihood analyses resulted in nearly identical topologies, showing that cave forms of *Trichomycterus* in Santander have evolved independently at least twice. One evolutionary event corresponds to *Trichomycterus rosablanca*, which was recovered with strong support (0.96 posterior probability, 72% maximum-likelihood bootstrap) as sister to *Eremophilus mutisii*, a species that lives in surface streams in the Bogotá River basin and has conspicuous eyes and shows profusely vermiculated dark coloration (Fig. 2, Clade 1). The second evolutionary event corresponded to a clade including the species *T. sandovali* and specimens from the Sogamoso River drainage, including surface populations and all the remaining cave species and populations analyzed (Fig. 2, Clade 2). Relationships among the clades including *T. rosablanca* and *E. mutisii, T. sandovali* and allies, and other species were not strongly supported, but the Shimodaira-Hasegawa test showed that unconstrained and constrained topologies differred significantly. The topology in which cave populations were monophyletic was less fit to the data than the unconstrained topology in which troglomorphic traits are homoplasious (P = 0.002). We thus reject the hypothesis of a single origin for the evolution of cave-living and associated phenotypes (i.e. loss of eyes and pigmentation).

**Figure 2.**
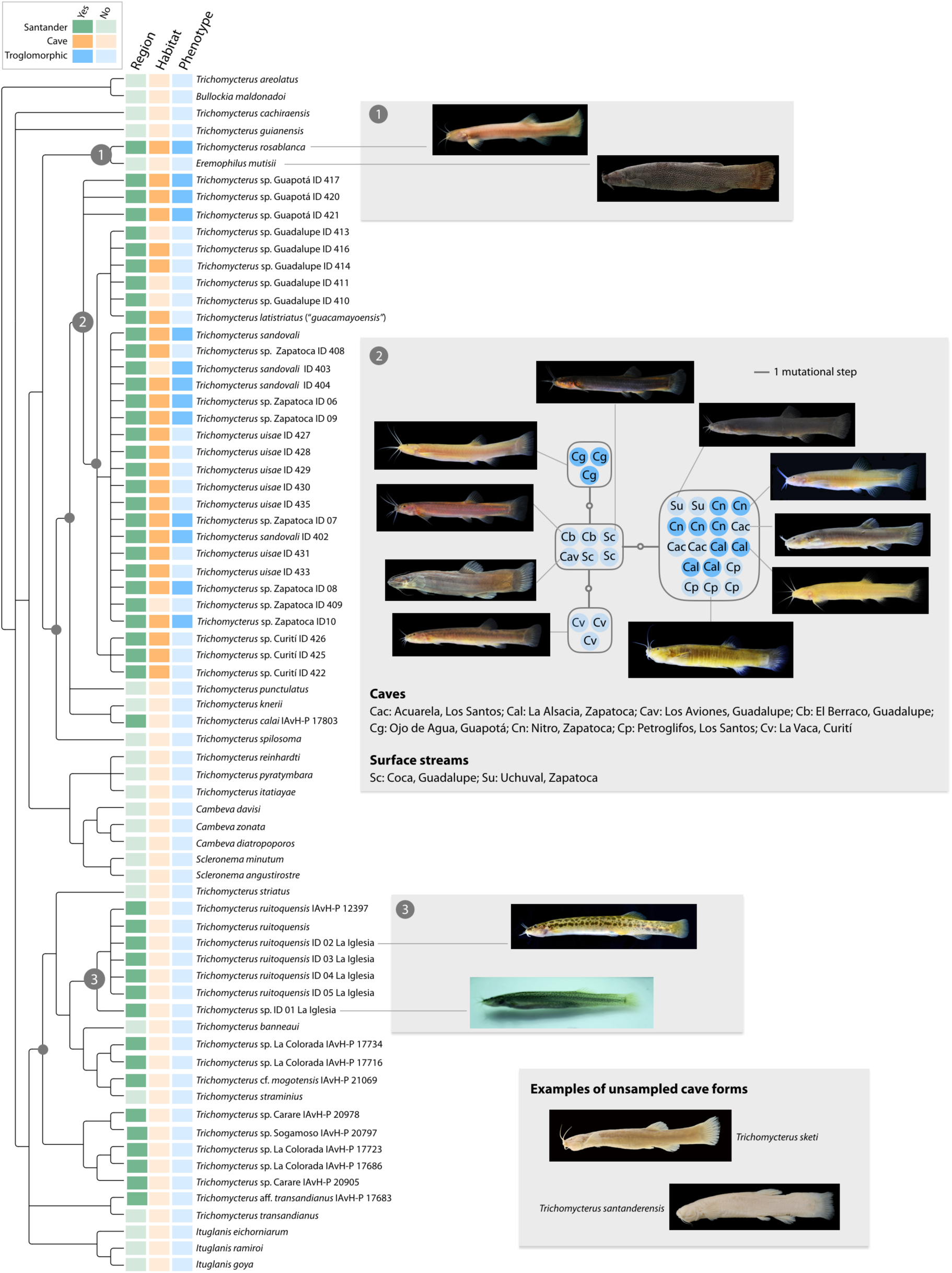
Relationships among catfish in *Trichomycterus* and other genera reveal repeated colonization of caves and phenotypic convergence in the karst region of Santander, Colombia. The Bayesian gene tree on the left is an overview of phylogeny based on sequences of the COI gene (outgroups not shown). Colors at the tips indicate whether specimens were collected in the western slope of the Eastern Andes of Santander, whether they were found in cave habitats, and whether they lacked eyes or body pigmentation (*i*.*e*. troglomorphism). Strongly supported nodes (posterior probability ≥0.95, maximum-likelihood bootstrap ≥70%) relevant to the phylogenetic position of specimens from Santander are indicated with circles; numbered nodes are mentioned in the text and in panels on the right. Node 1 defines a clade formed by phenotypically contrasting species from caves in Santander (*T. rosablanca*) and surface streams in the Bogotá River basin (*Eremophilus mutisii*). Node 2 defines a large clade for which genealogical relationships are shown on the right with a median-joining network. Rectangles correspond to haplotypes and circles to individuals, with colors indicating whether or not specimens lacked eyes or pigmentation following the color scheme in the phylogeny; upper-case letters indicate habitat (C= cave, S= surface) and lower-case letters define localities. Note that all specimens differ at most by two substitutions in COI, that individuals sharing two of the haplotypes occur in multiple localities including both caves and surface streams, and that haplotypes are not consistently associated with phenotypes. The most common haplotype occurred both in caves and surface streams, and was shared by individuals with various phenotypes and by populations assigned to at least three different species: *T. sandovali, T. uisae*, and “*T. guacamayoensis”* (*i*.*e. T. latistriatus*). Finally, node 3 defines a clade from Santander more closely allied to species from other regions in the Neotropics than to taxa from the karst region; photographs of live fishes illustrate differences in phenotype between specimens from the same surface stream which also differed genetically, highlighting a potentially undescribed species. The panel on the bottom illustrates two species from cave habitats in Santander lacking eyes and pigmentation which we were unable to sample and may represent other instances of convergence. Photographs by the authors and by Felipe Villegas (*T. rosablanca*), Cesar Castellanos (*T. santanderensis* and *T. sketi*), and from DoNascimiento *et al*. (2014; *E. mutisii*).

Not only did we find that cave populations referred to *Trichomycterus sandovali* are more closely allied to surface taxa than to the cave species *T. rosablanca*, but also that genetic divergence between cave-dwelling *T. sandovali* and other taxa from both surface streams and caves is shallow to non-existent. For example, we observed no genetic structure between cave and surface populations from Zapatoca: the surface population beared eyes and had a spotted to homogeneously dark coloration, whereas cave populations (one of them collected at the type locality of *T. sandovali*) had reduced eyes or lacked them and their coloration was faint. Moreover, there was no genetic structure between populations from Zapatoca and Los Santos (the latter collected close to the type locality of *T. uisae* Castellanos-Morales, 2008), which are separated by the deep Sogamoso River basin (Fig. 2; Clade 2). Likewise, cave and surface populations collected in Guadalupe and Guapotá (Suárez River drainage), and Curití (Fonce River drainage), differed only by two substitutions from those collected at the type locality of *T. sandovali*. The haplotype network for Clade 2 showed that cave and surface populations share haplotypes, suggesting recent divergence or gene flow (Fig. 2). Furthermore, we found no genetic differences in COI sequences between *T. sandovali* and two other cave species, namely “*T. guacamayoensis”* Ardila Rodríguez 2018 (recently shown to be a junior synonym of *T. latistriatus* (Eigenmann, 1917); see DoNascimiento & Prada-Pedreros, 2020) and *T. uisae* (Fig. 2).

Despite their geographic proximity to surface and cave populations from Santander, several surface taxa from our study area were more closely related to species from other regions (Fig. 2). The species *Trichomycterus ruitoquensis* Ardila Rodríguez, 2007, *T. straminius* (Eigenmann, 1917), and undescribed taxa recently collected in the Carare, Colorada, and Sogamoso rivers belonged to clades with species not occurring in Santander like *T. striatus* (Meek & Hildebrand, 1913) (restricted to Costa Rica and Panamá) and *T. banneaui* (Eigenmann, 1912) (from tributiaries in the middle Magdalena basin, Department of Tolima, Colombia).

We obtained two additional unexpected results. First, our data revealed a potentially undescribed species in the La Iglesia surface stream in the city of Bucaramanga, Santander. Most of the specimens we collected at this locality corresponded genetically and morphologically to *Trichomycterus ruitoquensis* (Fig. 2, Clade 1). However, one of the individuals collected in the same stream differed genetically from *T. ruitoquensis* by five substitutions in COI and morphologically by an irregular spotted pattern (*vs*. regular in *T. ruitoquensis*). Second, we observed atypical coloration among some surface individuals collected in the Uchuval stream, near the Nitro and Alsacia caves in Zapatoca. Some individuals showed a homogeneously dark pigmentation, which differed from other individuals from the same stream exhibiting a spotted pigmentation pattern on the body (see photographs in Fig. 2).

## DISCUSSION

Our study provides evidence for the repeated evolution of cave-living forms from surface ancestors in trichomycterid catfishes in a restricted geographic region in Colombia. At least two lineages (*Trichomycterus rosablanca, T. sandovali* and allies) colonized caves independently, and convergently evolved loss of eyes and body pigmentation in the karst region of Santander. These two lineages occur in separate drainages (Suárez and Carare rivers, respectively) not connected by river courses, further supporting the hypothesis that cave-living forms are derived from independent colonization events of subterranean environments in different areas. We lack DNA sequences from cave species of *Trichomycterus* from other countries in South America (Fig. 1A). However, the vast geographical distances separating them and overall congruence between geography and phylogeny in the group (Ochoa *et al*., 2017; Katz et al., 2018; Ochoa *et al*., 2020) suggests that species of *Trichomycterus* have independently colonized caves multiple times. In addition, it remains possible that cave species from Santander which we did not sample may have independently colonized these environments (Fig. 2). Examples include *T. santanderensis* Castellanos-Morales, 2007 from the Lebrija River drainage which we hypothesize belongs to the clade formed by *T. sandovali* and allies, and *T. sketi* Castellanos-Morales, 2011 which occurs in a separate drainage (Opón River) and may represent a distinct lineage.

Convergent evolution associated with cave living has been documented in other fish systems including *Astyanax mexicanus* (Dowling *et al*., 2002; Bradic *et al*., 2012), *Garra barreimiae* Fowler & Steinitz 1956 (Kirchner *et al*., 2017), and amblyopsids (Niemiller *et al*., 2012). In all cases, cave-dweller fishes exhibit similar phenotypic traits including absence -or reduction-of eyes and body pigmentation, suggesting adaptation to cave environments along similar trajectories. More broadly, such pattern speaks to the repeatability of evolution by natural selection owing to common selective pressures and potentially to shared genetic and developmental constraints (Losos, 2017).

In addition to our finding of independent colonization of caves in *Trichomycterus rosablanca* and *T. sandovali*, our work raises the intriguing possibility that different populations closely allied to the latter (*i*.*e*. members of Clade 2) may have independently colonized and adapted to caves in drainages along the Sogamoso drainage. Genetic similarity and haplotype sharing between surface and cave populations in this clade suggests that these populations diverged recently, that they still experience gene flow, or both. Gene flow in these karstic systems is likely because water flows in and in some cases out of caves, potentially allowing dispersal of adult fishes, larvae, or eggs. This result is consistent with studies on other cave organisms (*Astyanax mexicanus, Garra barreimiae*, and cave salamanders) in which cave and surface populations are also connected by gene flow (Bradic *et al*., 2012; Niemiller *et al*., 2008; Kirchner *et al*., 2017), possibly due to sporadic flooding. Given genetic exchange between surface and cave environments, selective pressures strong enough to counteract the effects of migration are needed to account for phenotypic evolution in cavefishes (Cartwright *et al*., 2017). Accordingly, some studies suggest that divergence in the face of gene flow is a main driver of the evolution of cave-dwellers (Juan *et al*., 2010; Bradic *et al*., 2012). Limited genetic divergence between surface and cave populations is also consistent with scenarios in which the origin of cave phenotypes is initially promoted by phenotypic plasticity (Rohner *et al*., 2013; Bilandžija *et al*., 2019) or via epigenetic mechanisms (Gore *et al*., 2018). Future work extending our analyses to genome-wide assays of genetic variation will allow us to better characterize gene flow, elucidate relationships and estimate timing of divergence among populations, and assess the genetic basis of phenotypes in the system.

In addition to suggesting that caves were likely colonized repeatedly and that penotypic evolution in *Trichomycterus* may occur rapidly and with gene flow, our finding of no genetic differentiation in the COI gene between cave-dweller and surface populations in the *T. sandovali* group (Fig. 2, Clade 2) may have taxonomic implications. In particular, we found no genetic structure among populations currently assigned to *T. sandovali*, “*T. guacamayoensis”* (*i*.*e. T. latistriatus*) and *T. uisae*, and specimens from Guadalupe, Guapotá and Zapatoca municipalities. Although the populations we sampled are separated by up to nearly a couple hundred of kilometers and by deep canyons, and although some of them lack eyes and pigments while others have these traits, they all are genetically too similar to distinguish them using the mitochondrial marker we employed, which is often used as a barcode for species identification (Hebert *et al*., 2003; Ward *et al*., 225, 2009). While lack of COI divergence may simply reflect recent diversification or gene flow, an alternative interpretation is that *T. sandovali* might not correspond to a cavefish species restricted to the Alsacia cave in Zapatoca as traditionally believed, but rather to a taxon with much broader distribution consisting of hypogean and epigean populations throughout the Sogamoso River basin, much like the case of the Mexican Tetra (Bradic *et al*., 2012). A taxonomic review with careful attention to phenotypic variation is needed to assess the status of populations in the group as separate species and to identify several specimens included in our analyses which may correspond to undescribed species.

The importance of more extensive sampling to better understand patterns of cave colonization and evolution in our study system is illustrated by comparing our results to those implied by previous phylogenetic studies of trichomycterids. Based on work showing that two cave species lacking eyes and pigmentation (*T. sandovali* and *T. rosablanca*) occupied similar phylogenetic positions in analyses with partly overlapping —and incomplete— sampling (Mesa S. *et al*., 2018; Ochoa *et al*. 2017; Ochoa *et al*. 2020), it appeared likely that cave species from Santander had a single origin. Our phylogeny was generally congruent with trees based on thousands of loci including less taxa (Ochoa *et al*. 2017; Ochoa *et al*. 2020), yet only by sampling multiple additional populations we were able to determine (1) that *T. sandovali* and *T. rosablanca* belong to different clades, implying separate colonization of caves; and (2) that more than one event of cave colonization may have occurred among *T. sandovali* and its close relatives. Because multiple cave populations exist in the Santander karst region and remain unstudied, additional work has potential to reveal other examples of convergence associated with cave living.

Our finding of individuals with homogeneously dark pigmentation differing from other individuals in the same surface stream exhibiting a spotted pigmentation pattern is intriguing and requires further study. A potential explanation is that homogeneously pigmented individuals might be the result of re-colonizations of the surface environment from caves. Because cave-dweller populations are uniformly depigmented, individuals recolonizing the surface may have recovered dark pigmentation yet not in a spotted pattern likely lost following cave invasion. Although this scenario is speculative and would need to be tested, we note that cave fishes may be common exceptions to Dollo’s law (Collin & Miglietta, 2008), which argues that once a complex trait is lost it is very unlikely to be re-acquired. Although cave-dwellers have been considered evolutionary “dead-ends” (Stern *et al*., 2017) owing to their specialization to underground life involving the loss of eyes and pigmentation, evidence from several lineages of organisms adapted to caves including fishes (Dillman *et al*., 2011), salamanders (Trajano & Cobolli, 2012), scorpions (Prendini *et al*., 2010), and amphipods (Copilaş-Ciocianu *et al*., 2018) indicates that reacquisition of surface phenotypes upon recolonization from caves is indeed possible.

To conclude, we have shown that trichomycterids from Santander have experienced convergent evolution as a likely consequence of repeated colonization of cave environments, and that such convergence has likely occurred in the face of gene flow and within a relatively restricted geographic area. More broadly, *Trichomycterus* from Santander have great potential for future studies addressing additional evolutionary, ecological, and developmental questions about processes leading to the extreme phenotypes that characterize cave-living. How does speciation occur in these organisms? What is the role of forces like selection, mutation, drift, and gene flow in trait evolution? What are the molecular, genetic, and developmental processes underlying the loss of complex organs such as eyes? How are cavefish populations genetically structured? What are the implications of population structure for their conservation? Our study has barely scratched the surface of what we anticipate to be a promising system to study these and many other questions.

## ACKNOWLEDGEMENTS

We are grateful to Yuranis Miranda, Alexandra Jiménez, Nataly Pimiento, Gerardo Bárcenas, Francisco Bautista, Diego González, and Cinthy Jimenez for help during field work; to Jorge Avendaño and Catalina Palacios for collaboration during lab work; and to Juan Gabriel Albornoz for support when visiting the IAVH-P collection. This project would have not been possible for JSF without the years of constant support of Graciela Flórez Villamizar; his efforts are dedicated to her. MT remembers that Javier Peña was the first to point to him those little white and eyeless fish when jointly exploring the Cueva del Nitro, Zapatoca, in the late 1990s. This research was supported by Fundación Iguaque, the Departamento de Ciencias Biológicas at Universidad de los Andes, and Santander Bio. Santander Bio was a project funded by the Sistema General de Regalías, administered by the Departamento Nacional de Planeación (BPIN 2017000100046), executed by the Gobernación de Santander, and operated by the Instituto de Investigación de Recursos Biológicos Alexander von Humboldt and the Universidad Industrial de Santander (Inter–administrative Agreement 2243, Gobernación de Santander). Specimens were collected under permits issued by ANLA (Resolution 004 of 2015) and by the Instituto de Investigación de Recursos Biológicos Alexander von Humboldt (Decree 1376 of 2013).

